# Thermophilic bacteria employ a contractile injection system in hot spring microbial mats

**DOI:** 10.1101/2025.08.11.669294

**Authors:** Vasil A. Gaisin, Corina Hadjicharalambous, Izabela Mujakić, Cristian Villena-Alemany, Jiangning Li, Michal Koblížek, Martin Pilhofer

## Abstract

Bacterial contractile injection systems (CISs) are multiprotein complexes that facilitate the bacterial response to environmental factors or interactions with other organisms. Multiple novel CISs have been characterised in laboratory bacterial cultures recently; however, studying CISs in the context of the native microbial community remains challenging. Here, we present an approach to characterise a new, bioinformatically predicted CIS by directly analysing bacterial cells from their natural environment. Using cryo-focused ion beam milling and cryo-electron tomography (cryoET) imaging, guided by 16S amplicon sequencing, we discovered that thermophilic *Chloroflexota* bacteria produce intracellular CIS particles in a natural microbial hot spring mat. We then found a niche-specific production of CIS in the structured microbial community using an approach combining shotgun metagenomics, proteomics, and immunogold staining. Bioinformatic analysis and imaging revealed CISs in other extremophilic *Chloroflexota* and *Deinococcota*. This *Chloroflexota*/*Deinococcota* CIS lineage shows phylogenetic and structural similarity to previously described cytoplasmic CIS from *Streptomyces* and probably shares the same cytoplasmic mode of action. Our integrated environmental cryoET approach is suitable for discovering and characterising novel macromolecular complexes in environmental samples.

## Introduction

Microorganisms drive many important biogeochemical processes on our planet. Their growth is determined by basic physicochemical conditions, but it also depends on interactions between different microbial species and between individual cells of a single species. These interactions are often facilitated by multiprotein complexes, such as archaeal and bacterial contractile injection systems (CISs) [1]. CISs show homology to contractile phage tails and act as nanoscale syringes injecting cargo molecules, often toxic effectors, into a target cell [2]. The architecture of diverse CISs is conserved and comprises a contractile sheath surrounding an inner tube, as well as a baseplate module. The triggered baseplate initiates CIS firing, i.e. the contraction of the sheath and expulsion of the inner tube [3]. Assembled CIS particles can have different cellular localisations, which dictate their mode of action. The type six secretion system (T6SS) is a CIS that injects effectors from the inside to the outside of the cells being anchored in the cytoplasmic membrane [4,5]. Extracellular CIS (eCIS) are found free-floating in the cytoplasm before being released to the extracellular space to target other cells [6–8]. The thylakoid-anchored CIS fires while being anchored in the cyanobacterial thylakoid membrane stacks [9]. The cytoplasmic CIS (cCIS) resembles eCIS; however, it is targeted to induce cell death in the CIS-producing cells upon stress, without being released into the environment [10–12].

The T6SS subtype 4, eCIS from *Algoriphagus machipongonensis*, and the thylakoid-anchored CIS belong to one phylogenetic clade and share clade-specific structural proteins despite their differences in mode of action [6]. Therefore, predictions of CIS mode of action can be unreliable when based only on sequencing data. However, when sequencing data and high-resolution *in situ* imaging of CISs are combined, the CIS mode of action can be identified more confidently. A key technology for imaging CIS in their cellular context in a near-native flash-frozen state is cryo-electron tomography (cryoET) [13]. CryoET has become a powerful tool for *in situ* imaging, resulting in high-resolution and three-dimensional reconstructions (cryo-tomograms) of cellular structure [14]. CryoET, however, is limited to relatively thin samples – up to approximately 500 nm of thickness. Hence, cryo-focused ion beam milling (cryoFIB) is often required to thin thicker samples and generate so-called lamellae that can be subsequently imaged by cryoET [14].

Previous studies of CISs have focused on model organisms that can be cultured in the lab. To better understand how bacterial CISs are employed in natural environments, we set out to investigate a novel CIS produced by a natural bacterial population. We chose a *Roseiflexus*- dominated microbial mat as a model system for three main reasons. First, bacteria of the genus *Roseiflexus* possess CIS-like genes in their genomes [15–17]. Second, *Roseiflexus* bacteria have been found in hot springs worldwide [18]. Third, *Roseiflexus* cells form a relatively large and prominent population in microbial mats [19–21]. Hot spring microbial mats have a stable multilayer organisation, which makes them a convenient model system for *in situ* studies of ecophysiological processes and microbial interactions that govern spatially structured microbial communities [19,22]. However, due to the complexity and size of the mats, we had to tackle significant challenges in making these environmental samples amenable to cryoET imaging and in integrating this technique with approaches from microbial ecology. Here we employed the combination of conventional ‘omics’ techniques with cryoFIB/ET and immuno-TEM, to investigate hot spring mats at the levels of microbial communities, cells, and macromolecular complexes, resulting in the discovery of previously uncharacterised CISs.

## Materials and Methods

### Bacterial strains and growth medium

*Roseiflexus castenholzii* DSM 13941^T^, *Calidithermus chliarophilus* DSM 9957^T^, and *Thermogemmatispora carboxidivorans* DSM 45816^T^ were obtained from DSMZ (German Collection of Microorganisms and Cell Cultures, Braunschweig). *Deinococcus grandis* CCM 3997^T^ and *Deinococcus aquatilis* CCM 7524^T^ were obtained from the Czech Collection of Microorganisms (Masaryk University, Brno, Czechia).

Cells of *R. castenholzii* were grown anaerobically and aerobically. Anaerobic culture was grown in DSMZ medium 87 (*Chloroflexus aggregans* medium) under illumination with a halogen lamp at 48 °C. Aerobic culture was grown in a modified PE medium [23] in an Erlenmeyer flask at 48 °C with shaking (200 rpm) in the dark. Composition of the PE medium (pH 7.5-7.7): 0.38 g l^-1^ KH_2_PO_4_, 0.39 g l^-1^ K_2_HPO_4_, 0.50 g l^-1^ (NH_4_)_2_SO_4_, 0.10 mg l^-1^ Vitamin B_12_ (V6629-100MG, Sigma-Aldrich), 5 ml l^-1^ basal salt solution, 10 ml l^-1^ BME vitamin mixture (B6891, Sigma-Aldrich), 4.00 g l^-1^ yeast extract. Composition of the basal salt solution: 24.65 g l^-1^ MgSO_4_·7H_2_O, 23.40 g l^-1^ NaCl, 2.94 g l^-1^ CaCl_2_·2H_2_O, 1.11 g l^-1^ FeSO_4_·7H_2_O, 84.10 mg l^-1^ MnSO_4_·H_2_O, 28.80 mg l^-1^ ZnSO_4_·7H_2_O, 29.20 mg l^-1^ Co(NO_3_)_2_·6H_2_O, 25.20 mg l^-1^ CuSO_4_·5H_2_O, 24.20 mg l^-1^ Na_2_MoO_4_·2H_2_O, 31.00 mg l^-1^ H_3_BO_3_, and 4.73 g l^-1^ EDTA disodium salt dihydrate.

Cells of *T. carboxidivorans* were grown in a modified DSMZ Medium 592 in an Erlenmeyer flask at 55 °C with shaking (200 rpm) in the dark. The medium composition (pH 6.0): 1.30 g l^-1^ (NH_4_)_2_SO_4_, 0.25 g l^-1^ MgSO_4_·7H_2_O, 0.28 g l^-1^ K_2_HPO_4_, 0.07 g l^-1^ CaCl_2_·2H_2_O, 0.02 g l^-1^ Fe Cl_3_·6H_2_O, 1 g l^-1^ yeast extract, 1 g l^-1^ tryptone, 1 ml l^-1^ SL-8 trace elements solution. Composition of SL-8 trace elements solution: 36 mg l^-1^ Na_2_MoO_4_·2H_2_O, 24 mg l^-1^ NiCl_2_·6H_2_O, 17 mg l^-1^ CuCl_2_·2H_2_O, 190 mg l^-1^ CoCl_2_·6H_2_O, 62 mg l^-1^ H_3_BO_3_, 100mg l^-1^ MnCl_2_·4H_2_O, 70 mg l^-1^ ZnCl_2_, 1.5 g l^-1^ FeCl_2_·4H_2_O, 5.2 g l^-1^ EDTA disodium salt dihydrate.

Cells of *C. chliarophilus* were grown in the DSMZ-recommended Castenholz medium in an Erlenmeyer flask at 50 °C with shaking (200 rpm) for 3 days. The medium composition (pH 8.0) was: 100 mg l^-1^ Nitrilotriacetic acid (Titriplex I), 60 mg l^-1^ CaSO_4_·2H_2_O, 100 mg l^-1^ MgSO_4_·7H_2_O, 8 mg l^-1^ NaCl, 103 mg l^-1^ KNO_3_, 689 mg l^-1^ NaNO_3_, 140 mg l^-1^ Na_2_HPO_4_·2H_2_O, 0.47 mg l^-1^ FeCl_3_·6H_2_O, 2.20 mg l^-1^ MnSO_4_·H_2_O, 0.50 mg l^-1^ ZnSO_4_·7H_2_O, 0.50 mg l^-1^ H_3_BO_3_, 25 μg l^-1^ CuSO_4_·5H_2_O, 25 μg l^-1^ Na_2_MoO_4_·2H_2_O, 46 μg l^-1^ CoCl_2_·6H_2_O, 1 g l^-1^ tryptone, 1 g l^-1^ yeast extract.

Cells of *D. grandis* were grown in PYE medium in an Erlenmeyer flask at 30 °C with shaking (200 rpm) in the dark. Composition of PYE medium (pH 7.2): 10 g l^-1^ yeast extract, 10 g l^-1^ peptone, 5 g l^-1^ NaCl.

Cells of *D. aquatilis* were grown in R2A medium (prepared according to manufacturer recommendation. NCM0188A, NEOGEN) in an Erlenmeyer flask at 30 °C with shaking (200 rpm) in the dark.

### Sampling hot spring mats

We organised two sampling campaigns to Rupite hot springs, Bulgaria (41.458310840335805, 23.2619133366458). During the first campaign on October 28^th^, 2021, we collected several cores from the local microbial mats (Fig. S1B, Supplementary material). The core samples were placed in cryovials and immediately stored in a dry shipper at 77 K. These snap-frozen samples were delivered frozen to the laboratory in Zurich and stored at -80 °C before subsequent experiments. Temperature and pH were measured using Multi 350i meter (WTW, Germany). Basic characteristics of the sampling sites are provided in table S1 (Supplementary material). During the second campaign on August 7^th^, 2023, we collected large fragments of the mats which were maintained at ambient temperature and delivered to the laboratory within 8 h and then kept at 46 °C with illumination by halogen lamp overnight before processing. Samples were collected at 15:00-16:00 local time (under clear sky conditions) in both sampling campaigns.

### DNA extraction

The samples were loaded into tubes with Lysis matrix A (MP Biomedicals) and disintegrated using FastPrep-24 5G (MP Biomedicals) equipped with QuickPrep Sample Holder (MP Biomedicals) with the following parameters: Speed=7.0 m/sec, time=45 sec, cycles=1. The homogenised samples were resuspended in 470 µl TE buffer (50mM Tris pH 8.0, 100 mM EDTA) supplied with 5 µl of 10 mg ml^-1^ RNAse A (10109169001, Sigma-Aldrich) and 25 μl of 10% sodium dodecyl sulfate. The tube was inverted 5-10 times by hand after each chemical was added to mix the solution. The mixture was incubated at 37 °C for 13 min. After incubation 7 μl of 18.5 mg ml^-1^ Proteinase K (03115887001, Sigma-Aldrich) were added, and then the mixture was incubated at 37 °C for 1 h. After incubation, 100 μl of 5M NaCl were added and mixed. Then 80 µl of the CTAB solution (10% Cetrimonium bromide, 0.7 M NaCl) were added. The mixture was incubated at 65 °C for 10 min. After the mixture cooled to room temperature, 700 µL of chloroform was added and mixed manually. The obtained emulsion was centrifuged at 20,000×g for 10 min at room temperature. The upper aqueous phase was transferred to a clean tube. The procedure was repeated with 600 µl of chloroform. Isopropanol (0.6V of the final volume) was added to precipitate DNA from the aqueous solution. DNA was pelleted by centrifugation at 10,000×g for 10 min at room temperature. The pellet was washed in cold 70% ethanol, dried at room temperature, and dissolved in 60 µl of EB buffer (10 mM Tris, pH 8.5). After incubation at 37 °C for 30 min and mixing, the DNA solution was stored at -20 °C.

### 16S rRNA gene amplicon analysis

The amplification of the V3-V4 region of 16S rRNA gene, library preparation and amplicon sequencing were done by Azenta Life Sciences (first sampling campaign) and Microsynth AG (second sampling campaign). Azenta Life Sciences employed proprietary PCR conditions and Illumina MiSeq platform in 2x250 bp paired-end configuration. Microsynth AG employed standard 16S rRNA gene primers 341F/805R, Illumina two-step PCR library, Illumina NovaSeq platform in 2x250 bp single-end configuration. Raw reads were quality-checked using FastQC v0.11.7 (Babraham Bioinformatics, Cambridge, UK). Subsequent analyses were done in R studio v3.6.1. The primer sequences were trimmed and read quality filtered using Cutadapt v1.16 maximum error (-e 0.1), quality cut-off (-q 20) and minimum length (-m 250) [24]. Reads were truncated using *filterAndTrim* (truncLen = c(225, 225), maxEE=c(2,5), truncQ=2) in the R/Bioconductor environment from DADA2 package v1.12.1 [25]. ASVs were constructed and chimeric sequences removed using the method “pooled”. Additional chimeric sequences were removed by selecting 370-430 bp length range. Taxonomic assignment of amplicons was performed using non-redundant SILVA r138.1 16S rRNA gene database using *dada2:assignTaxonomy*. To reduce singletons and doubletons, reads found less than 3 times in less than 20% of the samples were removed from subsequent analysis. Community composition barplots were generated using Phyloseq v1.30.0 [26] and ggplot2 v3.3.6 [27].

### Metagenomic analysis

Shotgun metagenomic sequencing of the mat DNA was done using NEBNext Ultra II FS DNA Library Prep Kit and Illumina NovaSeq S1 platform in 2x150 bp paired-end configuration. Raw reads from three metagenomes (green, yellow and red mat) were quality filtered and adapters removed using Trimmomatic v0.36 (LEADING:3 TRAILING:3 SLIDINGWINDOW:4:15 HEADCROP:9 MINLEN:100) [28] and reads were assembled using SPAdes v3.13.1 [29] (following kmers 21,33,55,77,127 --meta parameters). Obtained contigs (>2.5 kb) were used for binning with MetaBAT2 2.12.1 with default parameters [30]. CheckM [31] was used to estimate bin completeness and contamination. Poorly resolved bins with contamination >10% were further manually curated by removing contigs with different sequencing depths [32] which were calculated according to Aldeguer-Riquelme et al. 2023 [33] and with repeated rounds of contamination/completeness assessment by CheckM. Final curated bins with completeness >50% and contamination <10% were considered MAGs. They were further taxonomically classified using GTDB-tk [34] (Supplementary material, Table S2). In total 35 MAGs were assembled, 12 MAGs from the yellow layer, 9 from the green layer, and 14 MAGs from the red layer. Accession numbers of MAGs are provided in (Supplementary material, Table S2). Taxonomic annotation of the shotgun Illumina reads was done using kaiju with database kaiju_db_nr_2023-05-10 [35]

### Identification of sheath genes in metagenomic data and recruitment analysis

Two MAGs, one from green mat, and one from yellow mat were classified as *Roseiflexus* sp. Based on ANI/AAI analysis done using fastANI [36] and AAI: Average Amino acid Identity calculator [37] they represent the same species with 99.9 and 99.75% of identity, respectively. Both of the MAGs were annotated using Prokka [38] and analysed in Geneious Prime® 2023.2. to obtain the CIS cluster.

All three metagenomes were checked for the presence of any other sheath gene from non-redundant database consisting of 1,679 sheath gene sequences of various bacteria downloaded from eCIS database [15]. Fragment recruitment was used to assess abundances of MAGs in the metagenome. Metagenomic reads were mapped against the MAGs using BLASTn with a cutoff of 70% query coverage, *e*-value 0.1 and the ‘best hit’ option. Finally, abundances were calculated considering only the hits with identities ≥95%. To compare abundance across the distinct metagenomes, the number of recruited bases from the mapped reads was normalized by the total size of each metagenome (expressed as base pairs per megabase of the metagenome).

### Sample vitrification

Frozen mat samples from the first sampling campaign were thawed at room temperature. Small pieces were cut from the mat samples using a sterile scalpel and then softly homogenised into a cell suspension using a sterile glass mortar and pestle. The cell suspension was subsequently vitrified using a Vitrobot Mark IV System (Thermo Fisher Scientific Inc., USA) [39]. Briefly, we applied 4.5 and 3.5 μl of the cell suspension (first and second sampling campaigns, respectively) to glow-discharged copper electron microscopy (EM) grids (R2/2 and R2/1 H2, Quantifoil). The EM grids were then back-blotted for 5 or 9 seconds in the humidified (100%) Vitrobot chamber and plunged into liquid ethane-propane (37/63 Mol%/Mol%). Finally, the grids were stored in liquid nitrogen until further processing. The back-blotting was carried out using filer paper from the back of the grid and a Teflon sheet at the front.

### Cryo-focused ion beam milling

All lab cultures and the mat samples from the second sampling campaign were cryoFIB milled following the automated sequential FIB milling scheme previously described in [40]. A Crossbeam 550 FIB-SEM instrument (Carl Zeiss AG, Germany) was used, equipped with a copper-band cooled mechanical cryo-stage (Leica Microsystems GmbH) and an integrated VCT500 vacuum transfer system (Leica Microsystems). In short, frozen EM grids were loaded onto a pre-tilted cryo-holder in a VCM loading station (Leica Microsystems) and transferred to an ACE600 (Leica Microsystems) using the VCT500. The grids were coated with 4-7 nm of Tungsten before being inserted into the Crossbeam. In the Crossbeam, an additional layer of organoplatinum was applied. Scanning electron microscopy with the SE2 and in-lens secondary electron detectors (Carl Zeiss AG, Germany) was then used to determine grid quality (3-10 V, 58 pA) and to target cells (3 V, 58 pA). Using automated sequential FIB milling, lamellae of ∼250 nm thickness were generated at a milling angle of 12°-15°. Targets were sequentially milled using 700 pA, 300 pA and 100 pA for rough milling and 50 pA for polishing. Once milling was completed, the cryo-holder was transferred to the VCM loading station using the VCT500 where the grids were unloaded and stored in liquid nitrogen until further processing.

### CryoET data collection and processing

CryoET data were collected using a Titan Krios G4 cryo-electron microscope (Thermo Fisher Scientific Inc., USA) with an accelerating voltage of 300 kV. The microscope was equipped with a BioContinuum imaging filter and a K3 direct electron detector (Gatan). Tilt series were acquired using SerialEM [41,42] and PACE-tomo [43] following bidirectional or dose-symmetric tilt schemes with 2° or 3° increments at -8 μm defocus. For non-FIB-milled bacterial cells, the tilt series were collected over a range of −60° to +60° with a total electron dose of ∼159 e^-^/Å^2^ and pixel size of 4.51Å. To account for the pre-tilt, lamellae of cryoFIB-milled cells were imaged over a range of -50° to +70° with a total electron dose of 82-140 e^-^/Å^2^ and pixel size of 4.51Å.

The tilt series were drift-corrected using the alignframes command in IMOD [44]. Cryo-tomograms were reconstructed using IMOD with the back projection method and 4× binning. The tomograms were CTF-deconvolved and filtered using IsoNet to improve contrast [45]. Dragonfly was used to segment tomograms according to the published protocol [46]. UCSF ChimeraX was used to visualize the results of the segmentations [47].

### Isolation of CIS particles from bacterial cells and room temperature-TEM

Samples were placed into 2 ml tubes with Lysing Matrix A (116910050-CF, MP Biomedicals) supplied with 1 ml of cold sample buffer (150 mM NaCl, 50 mM Tris, pH 7.4, 2x cOmplete™, EDTA-free Protease Inhibitor Cocktail) and bead-beaten in FastPrep-24 5G (MP Biomedicals) using QuickPrep Sample Holder (MP Biomedicals) at 4 °C with the following parameters: Speed=7.0 m s^-1^, time=45 sec, cycles=1. The homogenate was mixed with lysis buffer (150 mM NaCl, 50 mM Tris, pH 7.4, 300 μg ml^-1^ lysozyme (62971-10G-F, Sigma-Aldrich), 60 μg ml^-1^ DNAse I (10104159001, Sigma-Aldrich), 0.5x Protease Inhibitor Cocktail) and incubated at 37 °C for 15 min. Cell debris was removed by centrifuging the lysed material at 15,000×g for 15 min at 4 °C. The CIS particles were pelleted by ultracentrifugation at 150,000 g for 1 h at 4 °C and resuspended in the cold sample buffer.

To image CIS particles, the preparations were applied to glow-discharged, formvar-coated copper grids (FCF200-CU 50/pk, Formvar/Carbon 200 Mesh) for 60 s, and stained twice with 1% phosphotungstic acid. The stained grids were analysed at room temperature using transmission electron microscope Morgagni 268 (Thermo Fisher Scientific Inc., USA) operated at 80 kV.

### Protein identification in the CIS preparations

Proteins in samples with isolated CIS particles were identified at the Functional Genomic Center Zurich. Briefly, proteins were precipitated with trichloroacetic acid (Sigma-Aldrich) at a final concentration of 5% and washed twice with ice-cold acetone. The pellet was air-dried and dissolved in a buffer (10 mM Tris, 2 mM CaCl_2_, pH 8.2). Then, proteins were reduced and alkylated by adding Tris(2-carboxyethyl)phosphine and 2-chloroacetamide to a final concentration of 5 mM and 15 mM, respectively. The samples were incubated for 30 min at 30 °C light protected. Then, samples were enzymatically digested with trypsin (pH 8) and dried. Peptides were acidified to perform a cleanup using home-made C18 stage-tips (peptides were loaded on the tip, washed, eluted, and dried). The digested samples were dissolved in aqueous 3% acetonitrile with 0.1% formic acid. Peptides were separated on a M-class UPLC and analysed on a Ion Trap mass spectrometer (Thermo Fisher Scientific Inc., USA).

### Immunogold labelling

5 µl of CIS preparations were applied on glow-discharged EM grids (FCF200-CU-50, Electron Microscopy Sciences) and incubated for 2 h at room temperature in a closed Petri dish with water in a smaller dish inside to prevent fast drying. The grid was washed with cold sample buffer and blocked with filtered 0.1% bovine serum albumin (A7030-50G, Sigma-Aldrich) for 1 h at room temperature. Then the grid was washed again and incubated overnight at 4 °C with custom polyclonal anti-sheath antibodies (diluted 1:50) or rabbit IgG control (A01008, GenScript). Then the grid was washed again and incubated for 2 h at room temperature with gold-conjugated anti-rabbit secondary antibodies (Sigma-Aldrich, code G7277-.4ML). Finally, the grid was washed, dried at room temperature, and stained with 1% phosphotungstic acid before being imaged on the TEM Morgagni 268 operated at 80 kV. All washing steps were done in a 1.5 ml tube with 500 μl of the cold sample buffer. The grid was blotted with blotting paper between washing and incubations. The tube was turned 5 times and incubated for 10-20 min at room temperature to complete washing. The anti-sheath antibodies were generated at GenScript (PolyExpress™ package SC1676) with target sequence: GKAPPPPRLELPTRASKALTSLIVTPKSETASDIQVEIGPPVGENPPPEAFTVKISMGEVKEVYENVSFNKRPKDGTSYVVEKINSSSTLVQVAEGPATGSLADRVPEFGMSVIKPLAPIVPARVDATTFVGSAAERS. The number of labelled CIS (sheath) particles and the total number of CIS particles were manually counted in images randomly collected from areas on the EM grids. We analysed 56 images for preparations from mat layers and 52 image for preparations from axenic cultures of *R. castenholzii* and *S. coelicolor*. Average number of the labelled CIS particles and average number of the total number of CIS particles were calculated by analysis of three independent mat cores. A CIS particle is classified as labelled if it is covered by at least four gold beads. One-way ANOVA followed by multiple comparison Šidák test was performed using GraphPad Prism version 10.5.0.

### Phylogenetic analysis of the CIS proteins

We used sequences of the baseplate (WP_012121309.1) and sheath (WP_012121318.1) proteins from *R. castenholzii* DSM13941 as query sequences for PSI-BLAST (blast.ncbi.nlm.nih.gov) search against all non-redundant GenBank CDS. Multiple protein sequence alignment was constructed using MAFFT with L-INS-I parameter (--inputorder -- maxiterate 1000 --retree 1 --localpair input) [48]. Phylogenetic tree was calculated using IQ-TREE 1.6.11 [49] with the LG+F+I+G4 model recommended by ModelFinder [50] and branch supports were estimated using UFBoot2 [51]. The calculated phylogenetic tree was visualized using iTOL [52].

## Results

### Rupite hot spring mats accommodate a substantial Roseiflexus population

We collected microbial mats from the Rupite hot spring, which is known to accommodate a population of *Roseiflexus* bacteria [53,54]. After the first sampling campaign, we extracted genomic DNA from the snap-frozen mat samples and conducted 16S amplicon sequencing, which confirmed the presence of a substantial population of *Roseiflexus* bacteria in the samples from the high-temperature sites (> 48 °C) of the hot spring (Fig. S1, Table S1, Supplementary material).

### Environmental cryoET reveals an intracellular CIS in Roseiflexus-like cells

As an initial step in our cryoET approach to environmental samples, we examined the condition of mat cells after recovery from their snap-frozen state. Recovered mat samples were gently homogenised, plunge-frozen and screened by cryoET. This revealed that cells had lost their integrity (Supplementary material, Fig. S2). Therefore, we collected new mat samples and delivered them from the sampling site (Fig. 1A) to our workflow without snap-freezing to minimise the loss of cellular integrity.

**Figure 1.**
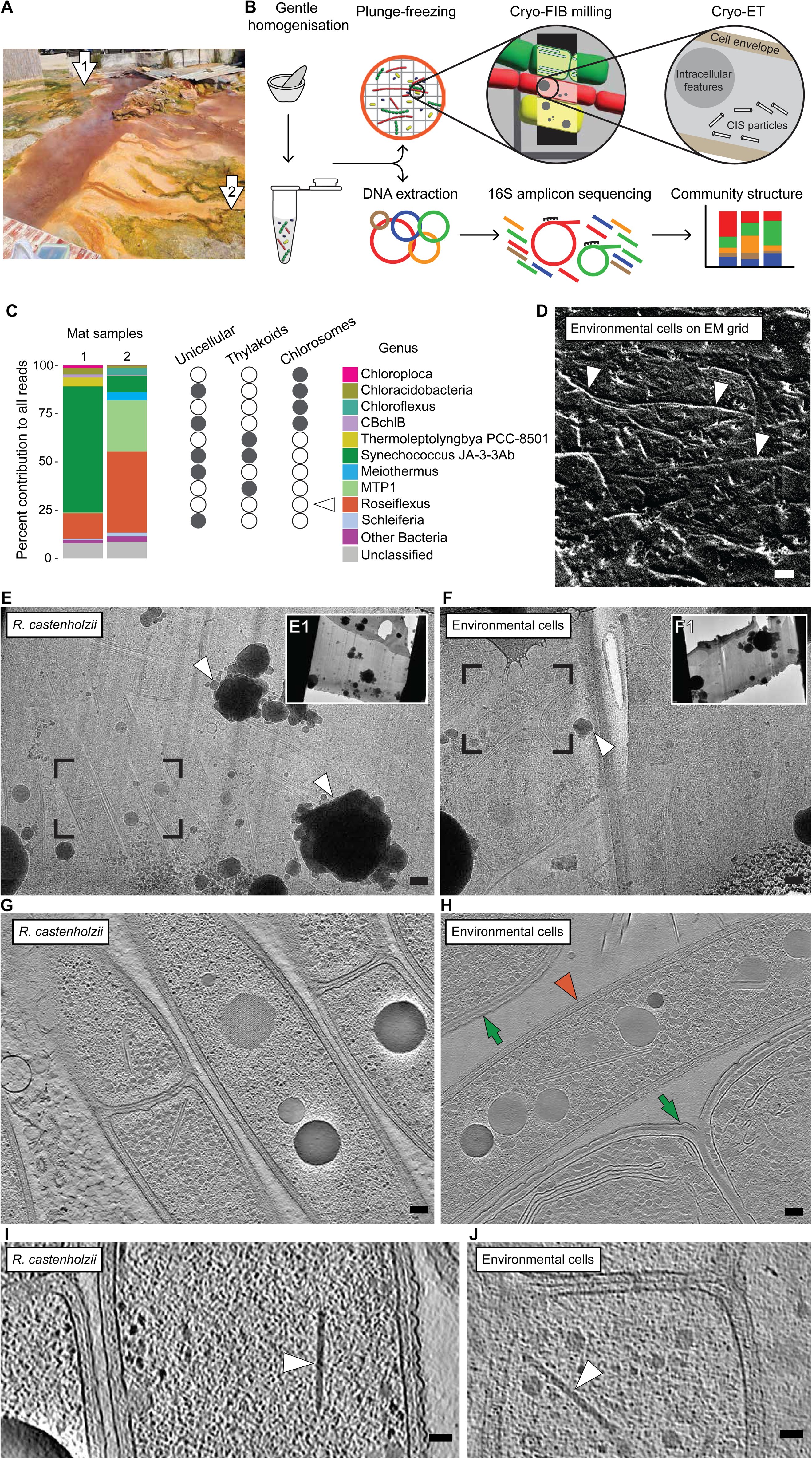
Environmental cryoET unveils intracellular CIS in the mat cells. (A) Rupite hot spring during second sampling campaign. Arrows point to the sampling site. (B) Scheme showing the workflow for environmental cryoET. (**C**)The bar plot shows the relative abundance of *Roseiflexus* among the top ten bacterial genera according to 16S amplicon sequencing results. The circles show presence (grey) and absence (white) of the morphological traits. Arrowhead points on traits of *Roseiflexus* bacteria. (**D**) CryoSEM image shows mat cells (arrowheads) on surface of EM grid. (**E**) Cryo-TEM image shows lamella milled through aerobic *R. castenholzii* cells. Full overview of lamella is shown in the right top incretion (E1). The box indicates the area used to collect cryoET data, which were then reconstructed into the cryo-tomogram shown in G. The arrowhead indicates ice contamination. (**F**) Cryo-TEM image shows milled lamella of mat homogenate on an EM grid. Full overview of lamella is shown in the right top incretion (F1). The box indicates the area used to collect cryoET data which were then reconstructed into the cryo-tomogram shown in H. The arrowhead indicates ice contamination. (**G**) Slice through a cryo-tomogram shows aerobic *R. castenholzii* cells. (**H**) Slice through cryo-tomogram shows cyanobacterial (arrows) and *Roseiflexus*-like cells (arrowhead) from the mat. **(I/J**) Slices through cryo-tomograms of *R. castenholzii* (**I**) and environmental *Roseiflexus*-like cells (**J**) show CIS-like intracellular features (arrowhead). Slice thickness: 18 (G, H, I) and 13.5 (J) nm. Scale bars: D 10 μm; E, F 300 nm; G, H 100 nm; I, J 50 nm.

We cut small fragments from the fresh mat samples, gently homogenised them to generate a cell suspension, and split the suspensions into aliquots for further cryoET analysis and DNA extraction (Fig. 1B). The genomic DNA was used to conduct 16S amplicon sequencing, which was an important step in our workflow, because it generated a list of the bacteria present in the mat samples. We found that within the top ten bacterial genera present in the mat, *Roseiflexus* bacteria were the only bacteria with known filamentous multicellular morphology and without thylakoids and chlorosomes (Fig. 1C). The genus *Roseiflexus* contributed 13% and 42% of the total reads in sample 1 and sample 2, respectively. Therefore, we assumed to find a high abundance of *Roseiflexus* cells in the mat material frozen on EM aliquoted for further cryoET imaging.

To acquire environmental cryoET data, we plunge-froze the mat cell suspensions on EM grids. Cryogenic scanning electron microscopy (cryoSEM) of the EM grids revealed a relatively thin layer of the mat cells (Fig. 1D), which was only achieved due to a homogenisation step prior to plunge-freezing. A relatively thin layer of the sample is essential for sample vitrification, since plunge-freezing is limited by sample thickness [55]. Additionally, the thin sample layer allowed for automated cryoFIB milling of the mat cells allowing us to generate 34 thin lamellae from the bulk material of the frozen mat cells.

Guided by the list of the bacterial species in the sample, we searched the *Roseiflexus*-like morphotypes in our images. In parallel, to improve the confidence in targeting *Roseiflexus* cells in the mat lamellae, we cryoFIB-milled lamellae of cells from axenic cultures of *R. castenholzii* (reference lamellae). Images of *R. castenholzii* cells in the reference lamellae (Fig. 1E) were used to identify *Roseiflexus*-like cells and select priority areas in the mat lamellae for data acquisition (Fig 1F). However, to better understand the cellular composition of the entire mat, we collected cryoET data in areas with and without potential *Roseiflexus* cells, resulting in 89 cryo-tomograms.

To confidently identify *Roseiflexus*-like cells in the cryo-tomograms, we collected reference cryo-tomograms on lamellae cryoFIB-milled through anaerobic (19 cryo-tomograms) and aerobic (71 cryo-tomograms) axenic culture of *R. castenholzii* (Fig. 1G). We then catalogued the cellular features of *R. castenholzii*. These features included regularly organised sheets, flat membranous intracellular features, vesiculations of the cell membrane, septal channels, and putative polyhydroxyalkanoate-, polyphosphate-, and glycogen granules (Supplementary material, Fig. S3). Organisation of the cell envelope, especially at the cell-cell junction, was also used as an important feature characterising *Roseiflexus* cells.

When examining the mat cryo-tomograms, we utilised the catalogue of *Roseiflexus* cellular features to confirm that the cells we targeted in the lamellae were indeed *Roseiflexus*-like cells (Fig. 1H). Overall, we found 33 environmental *Roseiflexus*-like cells in our mat cryo-tomograms.

While analysing cryo-tomograms of *R. castenholzii* cells and *Roseiflexus*-like cells from the mat, we found intracellular rod-like features (Fig. 1I/J). In both *R. castenholzii* and mat cells, the CIS-like particles were freely deposited in the cytoplasm without any anchoring to the cell envelope, resembling an extended form of eCIS/cCIS [6,10]. Interestingly, these CIS-like particles were absent in tomograms of *R. castenholzii* cells from anaerobic cultures (Supplementary material Fig. S4). The average length of the CIS-like particles was 217±16 nm (n=218) in aerobically grown *R. castenholzii* cells. In the mat dataset, we found up to twelve examples of the *Roseiflexus*-like cells with intracellular CIS-like particles (Fig. 1J, Supplementary material, Fig. S5). The average length of these particles was 251±26 nm (n=14).

To conclude, cryoFIB/cryoET techniques guided by 16S amplicon sequencing enabled us to visualize bacterial cells collected directly from the natural environment and discover a new eCIS/cCIS-like protein complex in these cells.

### Roseiflexus cells are a major contributor to the CIS pool of the microbial community

While we used cryoET to show the presence of CIS in environmental *Roseiflexus* cells *in situ*, we aimed to put the *Roseiflexus* CIS into the context of the whole *Roseiflexus* population and into the context of the microbial mat community. Therefore, we applied a holistic approach to the snap-frozen multi-layered mat samples (first sampling campaign) in order to analyse the distribution of putative CIS genes in the microbial community. The mat was dissected into red, green, and yellow layers (Fig. 2 A). We assumed that every layer represented a specific micro-niche in the mat community because every layer was dominated by differently pigmented microorganisms.

**Figure 2.**
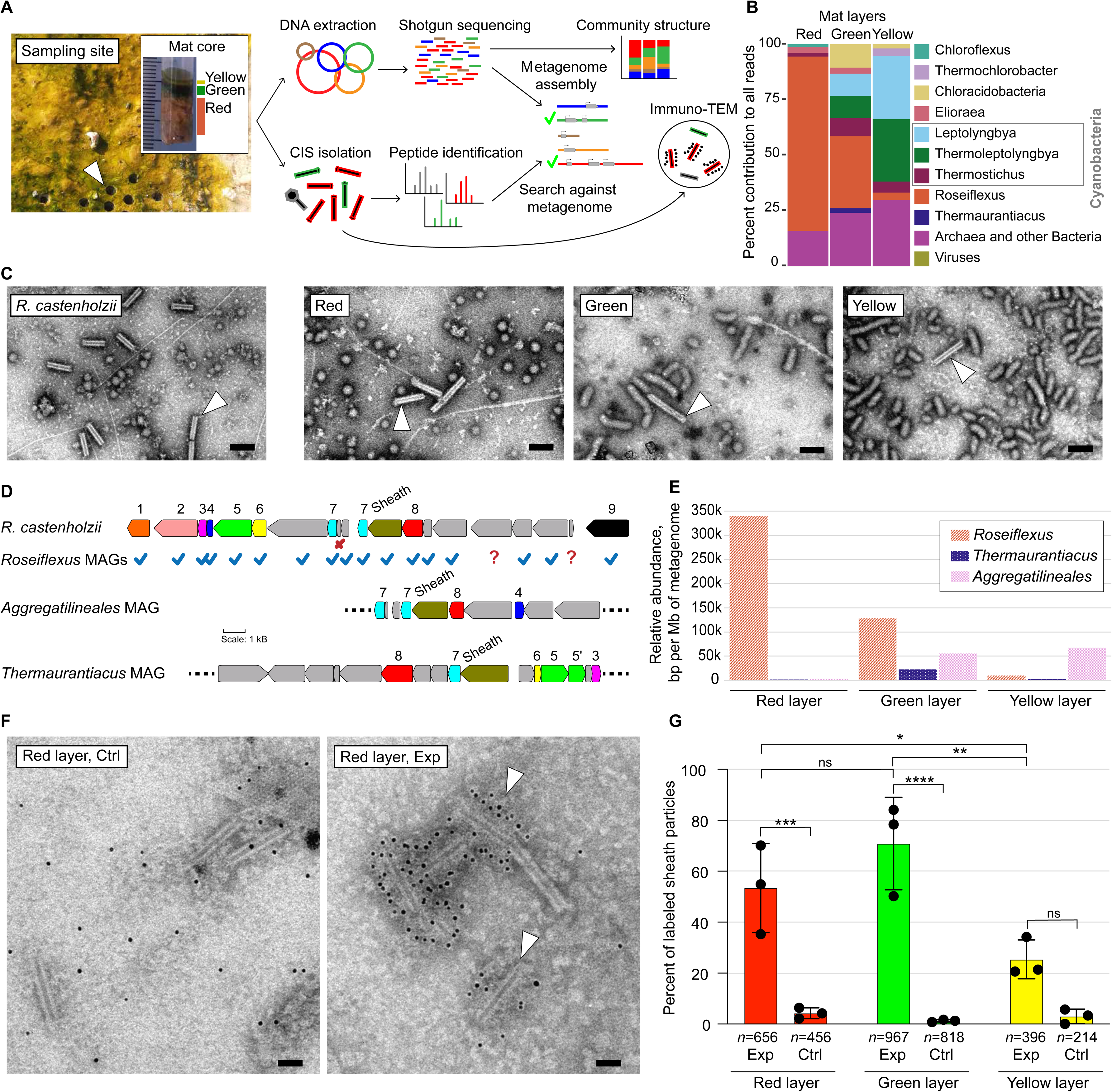
*Roseiflexus* population is a major contributor to the mat CIS pool. (**A**) Scheme showing the workflow from collecting mat cores (top left image) to metagenome assembly and immuno-TEM. (**B**) The bar plot shows the relative abundance of *Roseiflexus* among the top ten microbial species, based on the taxonomic annotation of shotgun Illumina reads using Kaiju. (**C**) TEM images show CIS (sheath) particles purified from an *R. castenholzii* axenic culture and each of the mat layers. Arrowhead points at the contracted CIS (sheath) particles. (**D**) The scheme shows the CIS gene cluster of *R. castenholzii* (used as reference) and the CIS gene locus in the Rupite MAGs (check mark = presence of a gene, x = absence of a gene,= gene not found). The numbers correspond to gene annotations: 1, 2, 3, putative baseplate genes; 4, 5, 6, putative spike genes; 7, inner tube gene; 8, putative apical cap gene; 9, ATPase gene. (**E**) The bar plot shows relative abundance of *Roseiflexus* sp., *Thermaurantiacus* sp., and *Aggregatilineales* bacterium from the different mat layers calculated based on normalized read mapping (the number of base pairs [bp] per megabase [Mb] of metagenome). (**F**) TEM images show examples of labelled CIS particles (arrowhead) from the mat samples. (**G**) Bar plot showing the distribution of the labelled CIS particles in the mat samples. Primary anti-sheath rabbit antibodies were applied for labelling the sheath (Exp). Rabbit IgG control was applied as control to check the unspecific binding of anti- rabbit gold-conjugated secondary antibodies (Ctrl). Here, *n* represents the total number of CIS (sheath) particles counted across the three experiments. Statistical significance between the samples was assessed using ordinary one-way ANOVA followed by multiple comparison Šidák test. Multiplicity adjusted P value was reported [ns (not significant), 0.1234; *, 0.0332; **, 0.0021; ***, 0.0002; ****, <0.0001]. Scale bars: C, 100 nm; F, 50 nm.

To identify all potential CIS-producers in the mat, we conducted shotgun Illumina sequencing of DNA from the mat layers and discovered all bacteria with CIS genes in the microbial community (Fig. 2A). Taxonomic assignment of the reads showed that *Roseiflexus* and cyanobacteria were the dominating bacterial genera (Fig. 2 B). We assembled, translated and annotated the metagenome from our shotgun DNA sequencing dataset, resulting in the metagenome of the mat and a set of metagenome-assembled genomes (MAGs) with completeness >50% and contamination <10% (Supplementary material, Table S2). Interestingly, *Roseiflexus* MAGs were assembled from the green and yellow layers but not from the red layer, even though *Roseiflexus* sp. is the most abundant bacterium in this layer. This occurred probably due to the high strain diversity of the *Roseiflexus* population within the red layer. The high microdiversity of co-occuring abundant microbial species is likely to generate assembly problems and reducing the quality of metagenome assembly [56].

Next, we lysed the mat cells and pelleted all large macromolecular complexes from the layers to purify CIS particles. Again, as for environmental cryoET, we used an axenic aerobic culture of *R. castenholzii* as a reference sample. We found similar empty CIS-like particles (contracted sheath) in preparations from the reference sample and from each of the mat layers (Fig. 2 C). To identify the proteins in these preparations, we performed mass spectrometry with peptide search against all our metagenomic contigs longer than 1 kb. The search resulted in 182 proteins being identified in the mat layers with 68 proteins being annotated as hypothetical proteins (Supplementary material, Fig. S6, Table S3). Using a BLASTp search against GenBank, we found sheath proteins among those hypothetical proteins. Additionally, we found cyanophage-related proteins belonging to *Elainellaceae* MAG (former *Leptolyngbya* sp. O-77) (Supplementary material, Table S4).

The sheath genes were found in the contigs that belonged to *Roseiflexus* sp., *Thermaurantiacus* sp., and *Aggregatilineales* sp. MAGs. CIS proteins are typically encoded by genes that are organized as a gene cluster on the bacterial chromosome or plasmid [15]. Using BLASTn search, we found other CIS-related genes down- and upstream of the sheath genes and assigned this locus as the *Roseiflexus* CIS gene cluster (Fig. 2 D). We also found CIS gene clusters in *Thermaurantiacus* and *Aggregatilineales* MAGs (Fig. 2 D). The *Roseiflexus* MAGs comprised all structural CIS genes predicted previously in the genome of *R. castenholzii* [15,16], but dispersed in multiple contigs in the MAGs (Fig. 2 D). We mapped the metagenomic reads to the three MAGs and found that *Roseiflexus* was the most abundant of these organisms and potentially the major contributor to the CIS pool in the red and green layers (Fig. 2 E).

To confirm that *Roseiflexus* is the major contributor to the mat CIS pool, we developed an immuno-TEM protocol and designed anti-sheath antibodies for the identification and quantification of *Roseiflexus* CIS particles. To design the antibodies, we aligned the sheath protein sequences from *Roseiflexus* genomes with the sheath sequence from *S. coeilicolor* to find domain 3, which forms the outer surface of the contracted sheath from *Streptomyces* cells [10]. The selected peptide sequence was used for generation of the anti-sheath rabbit antibodies.

The anti-sheath antibodies were applied as primary antibodies to the CIS preparations from *R. castenholzii* and *S. coeilicolor*, which we considered as positive and negative controls, respectively. Anti-rabbit gold-conjugated secondary antibodies were applied to the CIS preparations after incubation with anti-sheath antibodies and resulted in labelling of 99.2 % and 0.3 % of particles in positive and negative controls, respectively (Supplementary material Fig. S7). Also, we validated a relatively low rate of unspecific binding by secondary antibodies in the absence of the primary antibodies but in the presence of rabbit IgG control (Supplementary material Fig. S7).

The anti-sheath antibodies labelled CIS (sheath) particles in preparations from the mat layers (Fig. 2 F, Supplementary material Fig. S8). 53.4% and 70.9% of the particles were labelled in the red and green layers, respectively (Fig. 2 G). The yellow layer had only 25.4% of the particles labelled. These average numbers were calculated by analysis of three independent mat cores (Supplementary material Fig. S9).

Despite the red layer containing at least twice the *Roseiflexus* population of the green layer, CIS particles from *Roseiflexus* cells comprise an equally large fraction of the CIS pool in the green layer as in the red layer. These results indicate that *Roseiflexus* cells in the green layer produce more CIS particles than those in the red layer. Thereby, combination of DNA sequencing method, protein mass spectrometry and immuno-TEM enabled us to discover a niche-specific expression of *Roseiflexus* CISs in the bacterial populations in their natural environment.

### The Roseiflexus CIS belongs to a clade of cytoplasmic CISs from extremophilic bacteria

In the presented work, *Roseiflexus* bacteria were the first confirmed CIS-producing thermophilic bacteria. Previously, hypothetical CISs from *R. castenholii* were placed into CIS lineage IId (Fig. 3 A) together with cytoplasmic CISs from *Streptomyces* [15]. To characterise this system further, we searched for CIS gene clusters in other bacteria using the *Roseiflexus* sheath gene as a query for Position-Specific Iterated BLAST against the GenBank database (non-redundant protein sequences). We found new CIS gene clusters in multiple genomes of bacteria belonging to *Chloroflexota* and *Deinococcota* phyla (Fig. 3 B). Next, we reconstructed a maximum likelihood phylogenetic tree based on the concatenated sheath and baseplate J-like protein (Afp11) proteins and found that these CISs indeed form a clade outside cytoplasmic CISs (*Streptomyces*) but within the clade IId (Fig. 3 B). Interestingly, CISs from thermophilic *Deinococcota* strains clustered with *Chloroflexota* CISs but not with mesophilic *Deinococcota*. Additionally, the gene cluster organisation had higher similarity between *Chloroflexota* and thermophilic *Deinococcota* than between thermophilic and mesophilic *Deinococcota* (Fig. 3 C). These results suggest an acquisition of the CIS gene cluster via horizontal gene transfer between *Streptomyces*, *Chloroflexota* and *Deinococcota*.

**Figure 3.**
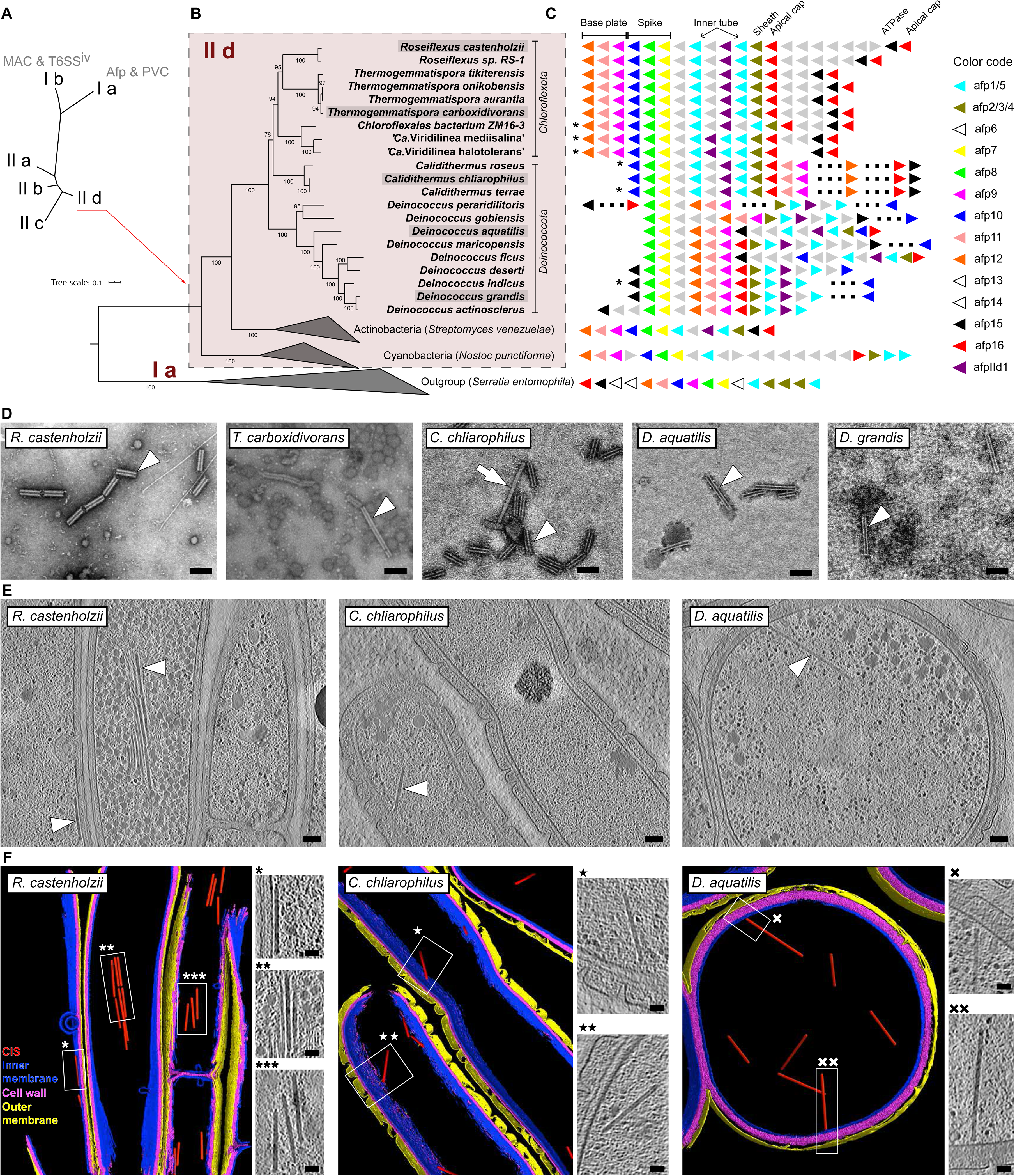
*Roseiflexus* CISs belong to a clade of cytoplasmic CISs from extremophilic bacteria. (**A**) Simplified scheme of the phylogenetic tree reconstructed by Chen *at al*. [10]. The scheme shows eCIS lineages. (**B**) Maximum likelihood phylogenetic tree showing a clade of *Roseiflexus*-related CISs. Version of the tree with expanded clades is shown in supplementary material (Fig. S10). The tree was reconstructed based on analysis of concatenated sheath and baseplate proteins. Grey highlighted strains were used for CIS preparations (shown in D). (**C**) Scheme shows an organisation of the CIS gene clusters in genomes from the *Roseiflexus* CIS clade, other members of CIS lineage subtype IId. Gene names were taken from Chen *at al.* [10]. Asterisk highlights draft genomes. Dashed line shows gaps between contigs. (**D**) TEM images showing CIS (sheath) particles purified from cells of the selected strains. Arrowhead points to the CIS particles. (**E/F**) Slices through cryo-tomograms (E) and corresponding models (F) showing the CIS particles (arrowhead/box) in cells of *R. castenholzii*, *C. chliarophilus*, *D. aquatilis*. Scale bars: D, E 100 nm; F (insertions), 50 nm.

To validate assembling CIS in *T. carboxidivorans, C. chliarophilus, D. aquatilis,* and *D. grandis*, we isolated CIS particles from their respective axenic cultures. *T. carboxidivorans* is a thermophilic mycelial *Chloroflexota* species [57]. *C. chliarophilus* is a thermophilic *Deinococcota* species [58,59]. *D. aquatilis* and *D. grandis* are mesophilic but UV/Gamma radiation-resistant *Deinococcota* species [58,59]. All preparations retrieved contracted sheath with rare examples of extended CIS particles as we have previously observed in preparations from *R. castenholzii* (Fig. 3 D). Next, we cryoFIB-milled the plunge-frozen cells of *C. chliarophilus* and *D. aquatilis* and imaged them with cryoET as representative examples of thermophilic and mesophilic *Deinococcota* in the CIS clade. In cryo-tomograms of these cells, we observed intracellular CIS particles (Fig. 3 E, F). The *Deinococcota* CIS particles were localised unattached in the cytoplasm, similar to CIS localisation in *Roseiflexus* cells. The average length of the intracellular CIS particles was 238±14 (n=69) and 365±39 (n=20) for *C. chliarophilus* and *D. aquatilis*, respectively. In conclusion, we discovered that *Roseiflexus* CIS is a member of new CIS lineage that is expressed and assembled in extremophilic bacteria.

## Discussion

Multiple novel CIS have been discovered in bacteria and archaea by studying microbial cultures in recent years. However, it was often difficult to place these systems back into the context of their native microbial environment. In this work, we looked at the CIS discovery from a different angle using an approach based on a combination of conventional omics techniques with cryoFIB/cryoET to analyse microbial cells from their natural habitat. Although we targeted specific bacterial cells and relied on the reference cryoET data, our study shows that cryoET can be applied as a discovery tool in environmental microbiology.

Since we validated the CIS assembly in environmental *Roseiflexus* cells, we can conclude that this system plays a role in the lifestyle of this bacterium under natural conditions. Phylogenetically, *Roseiflexus* CIS belongs to the same lineage (subtype IId [15]) as the recently discovered cytoplasmic CIS in *Streptomyces*, which has a cytoplasmic mode of action and induces the cell lysis under stress conditions [10–12,60]. Also, the CIS gene cluster in *Roseiflexus* genomes shares similarity regarding synteny with the CIS gene cluster in *Streptomyces* strains with validated function and was probably acquired via horizontal gene transfer between *Chloroflexota* and *Streptomyces*. Therefore, CIS in *Roseiflexus* bacteria may inherit the cytoplasmic mode of action. This hypothesis is further supported by the unanchored intracellular localisation of the CIS particles in *Roseiflexus* cells. Unfortunately, no genetic system is currently available for *Roseiflexus* strains, hence the involvement of *Roseiflexus* CISs in cell death and life cycle regulation remains unvalidated.

By studying cells directly collected from their natural environment, we discovered a niche-specific level of the CIS particle production. The high CIS production in the green layer may be linked to the cyanobacterial activity in this fraction of the mat community. The multi-layered organisation of the hot spring mats is thought to be formed as an adaptation of phototrophs (*Roseiflexus, Chloroflexus*, and Cyanobacteria) to utilise different wavelengths of light and hence reduce competition for the energy source [19]. The niche-specific CIS production lets us consider a new element of intercellular interactions governing this spatially structured microbial community. However, it remains unclear if *Roseiflexus* CIS is targeted towards cyanobacteria in eCIS-like manner or whether it acts as a self-targeted system controlling the *Roseiflexus* population in response to niche-specific conditions.

## Data availability

Example tomograms (XXX) were uploaded to the Electron Microscopy Data Bank.

Sequence data for all metagenomes generated in this work are deposited at EBI European Nucleotide Archive under the BioProject accession numbers PRJNA1041075 (yellow layer), PRJNA1041076 (green layer), PRJNA1041077 (red layer). The 16S amplicon sequencing data are deposited at EBI European Nucleotide Archive under the BioProject accession numbers PRJNA1307958.

## Acknowledgments

ScopeM is acknowledged for instrument access at ETH Zürich. Functional Genomics Center Zürich is acknowledged for mass spectrometry services and support. Authors acknowledges Dr. David Kaftan and Jason Lawrence Dean at Institute of Microbiology of the Czech Academy of Sciences for organising the first sampling campaign.

## Funding

MP was supported by the European Research Council (CoG 101000232) and the NOMIS Foundation. IM, CVA, and MK were supported by the OP JAK project Photomachines (reg. no. CZ.02.01.01/00/22_008/0004624) financed by the Czech Ministry of Education, Youth and Sports.

## Author contributions

VAG and CH collected samples, propagated bacterial cultures, and conducted microscopic analysis. IM and CVA analysed the DNA sequencing data. JL optimised culture of *R. castenholzii*. VAG, CH, and MP wrote the manuscript with input from IM, CVA, and MK.

## Competing interests

The authors declare no competing interests.

## Notes

### Competing Interest Statement

The authors have declared no competing interest.

### Summary of Updates

In this revision, we included an additional reference (Menzel et al., 2016, Nat. Commun.) and accession number for the 16S amplicon sequencing reads submitted to the EBI European Nucleotide Archive. We also added labels to Figure 1A and corrected Figure 2G and Supplementary Figure S7B.

