## Supplementary material for "Thermophilic bacteria employ a contractile injection system in hot spring microbial mats"

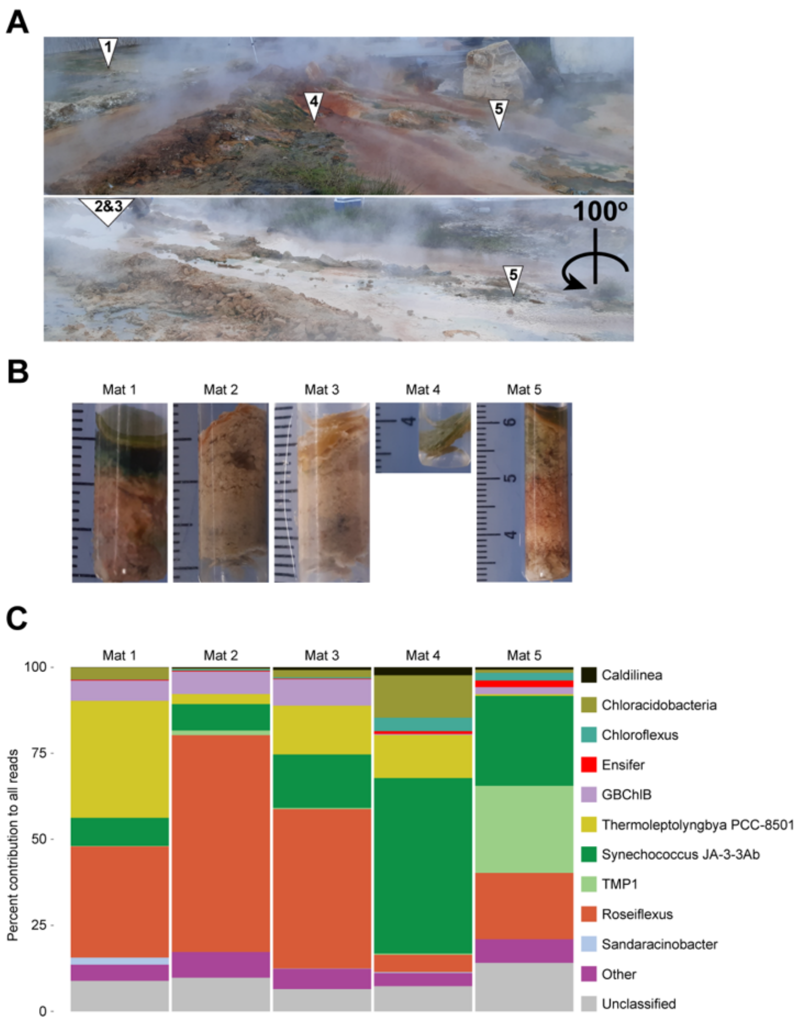


**Figure S1. Rupite hot spring accommodates a substantial *Roseiflexus* population. (A)** Picture shows the sampling sites at Rupite hot spring from two points of view. (**B**) Pictures show the mat cores collected at the corresponding sampling sites numbered in A. (**C**) Bar plot shows bacterial diversity in the mat samples based on 16S amplicon sequencing results.

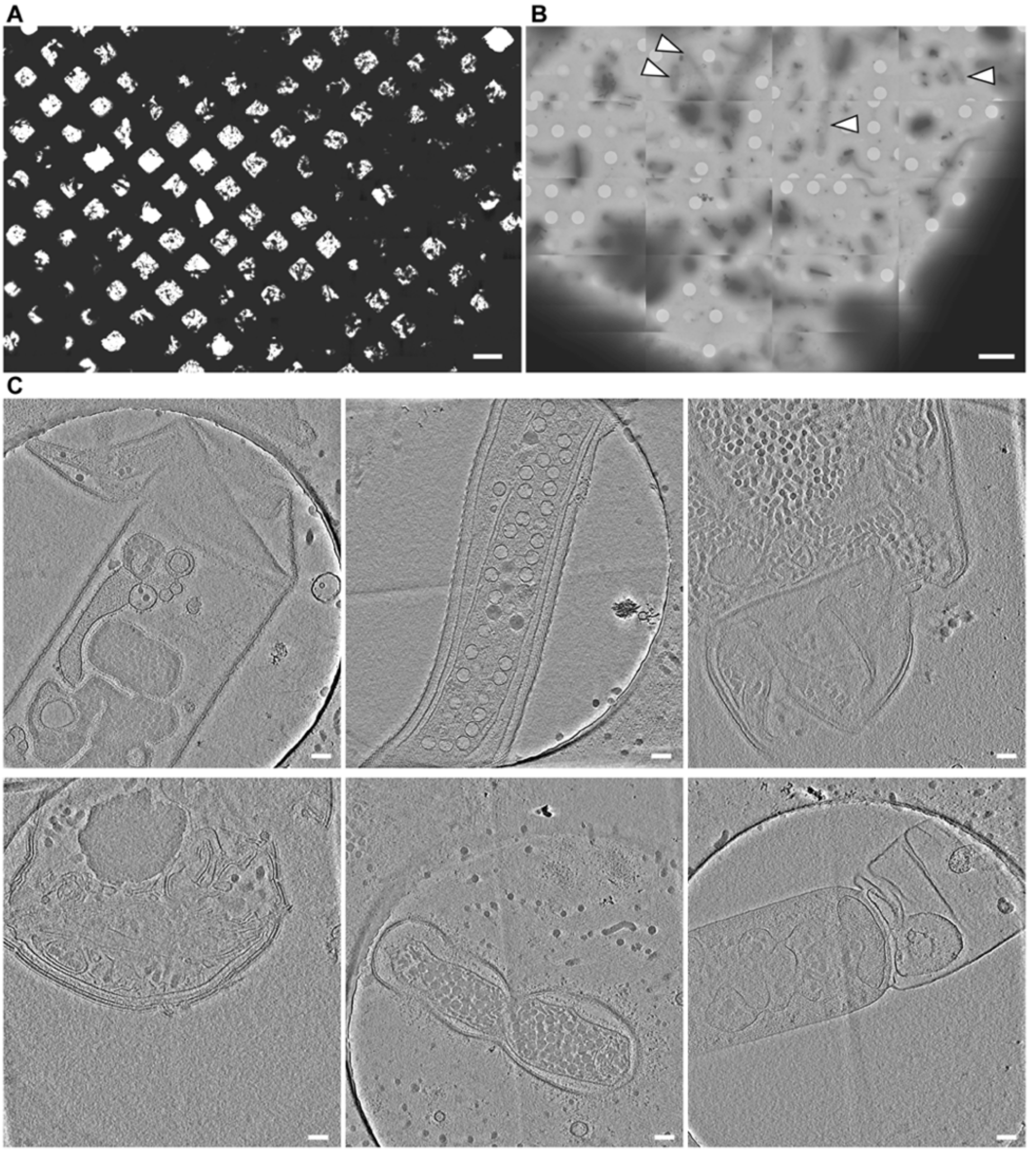


**Figure S2. Freeze-thaw cycles damage mat cells. (A)** The plunge-frozen mat material is shown in the overview image of an entire EM grid (magnification 135x). (**B**) The damaged cells are seen as "shadow" cells (arrowhead) in an overview image of a grid square (magnification 2,250x). (**C**) Examples of the damaged cells are shown in slices (18 nm thickness) through cryo-tomograms (magnification 19,500x). Scale bars: A, 100 μm; B, 5 μm; C, 100 nm.

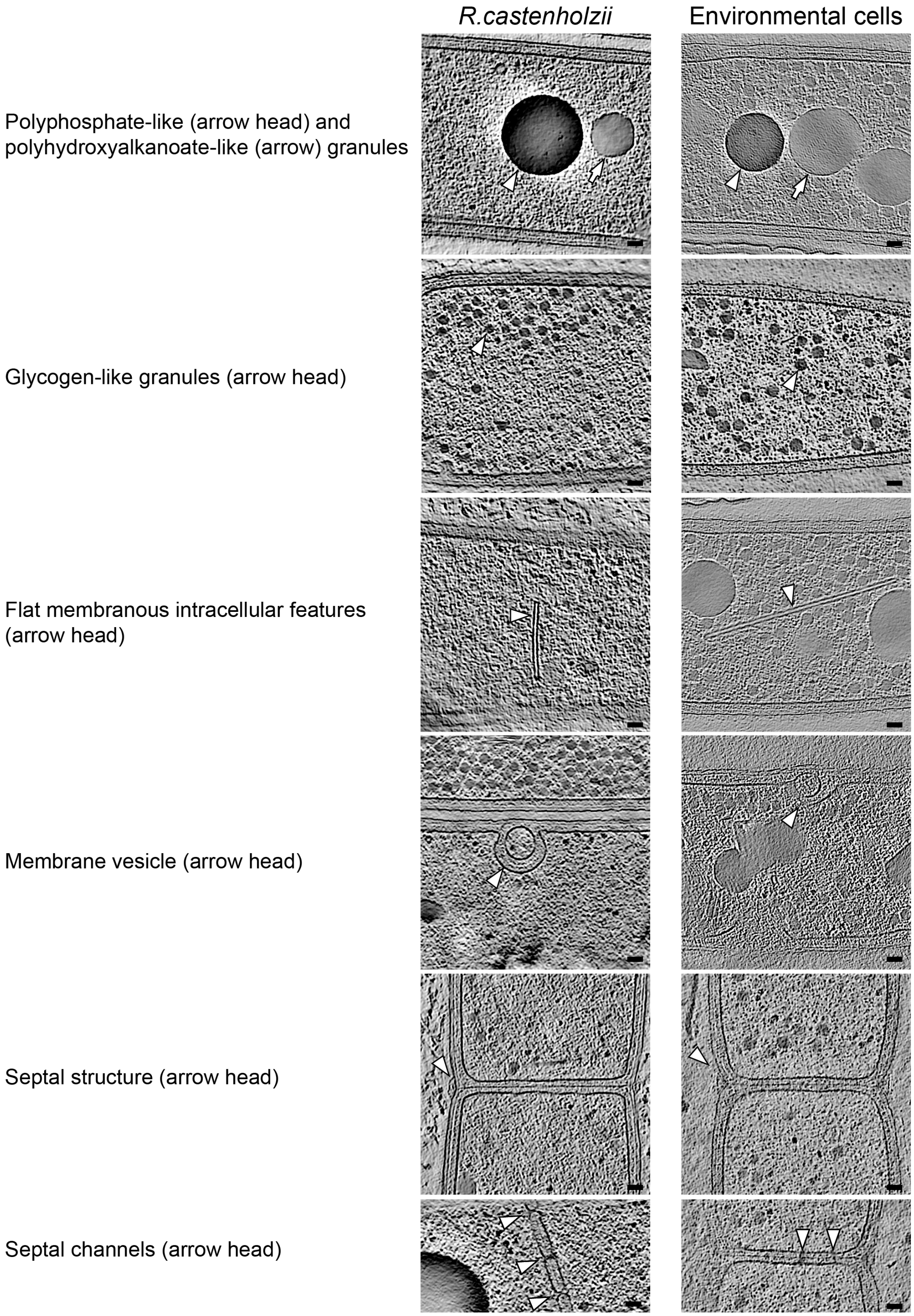


**Figure S3. *Roseiflexus* cells in axenic aerobic cultures and mats share unique cellular features.** Slices through cryo-tomograms show examples of cellular features in cells of *R. castenholzii* and in environmental *Roseiflexus*-like cells from the mat. Slice thickness: 18 nm. Scale bars: 50 nm.

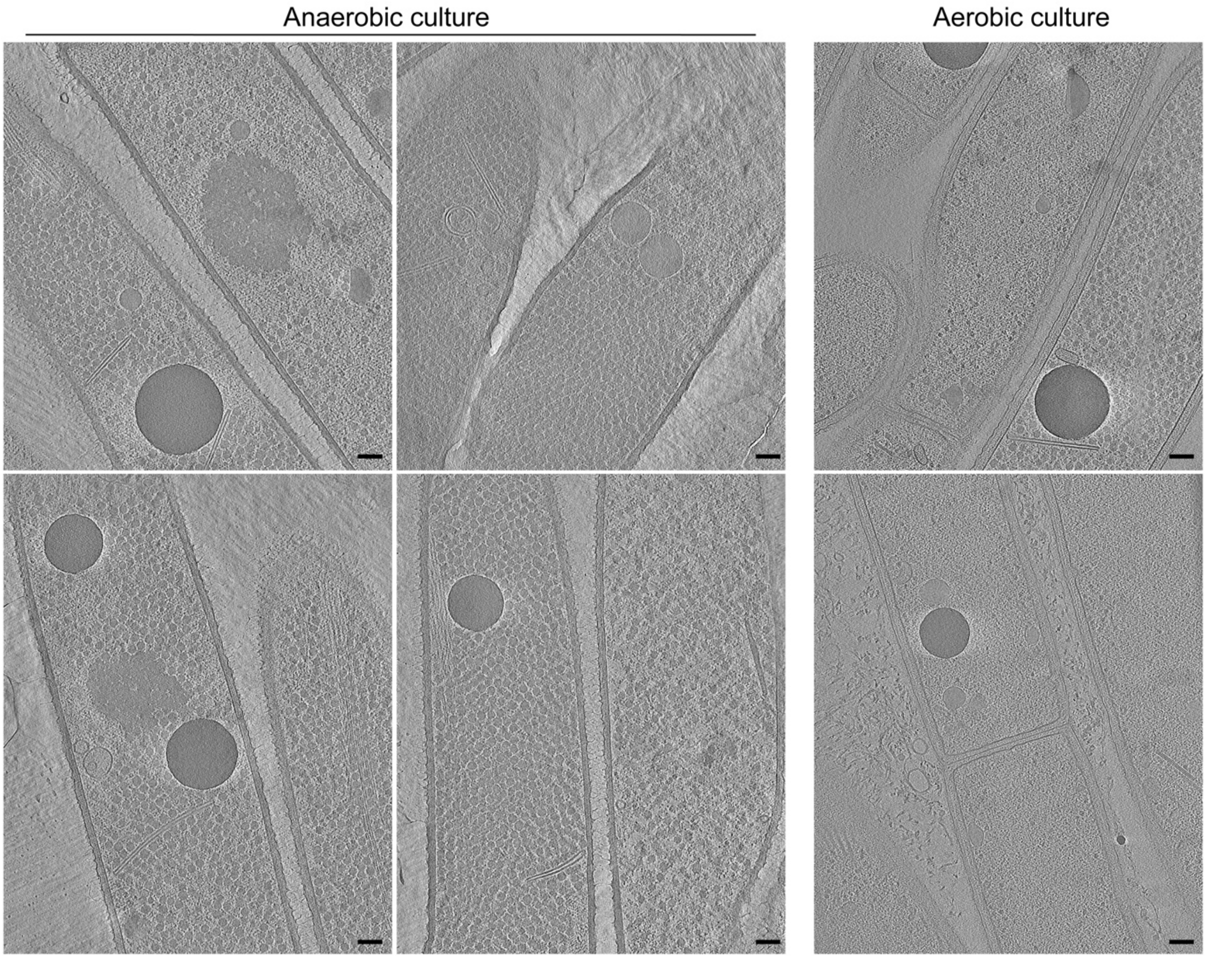


**Figure S4. Examples of cryo-tomograms with cells of *R. castenholzii* from the anaerobic and aerobic culture.** Slices through cryo-tomograms show examples of *R. castenholzii* cells from anaerobic and aerobic culture. Slice thickness: 18 nm. Scale bars: 100 nm.

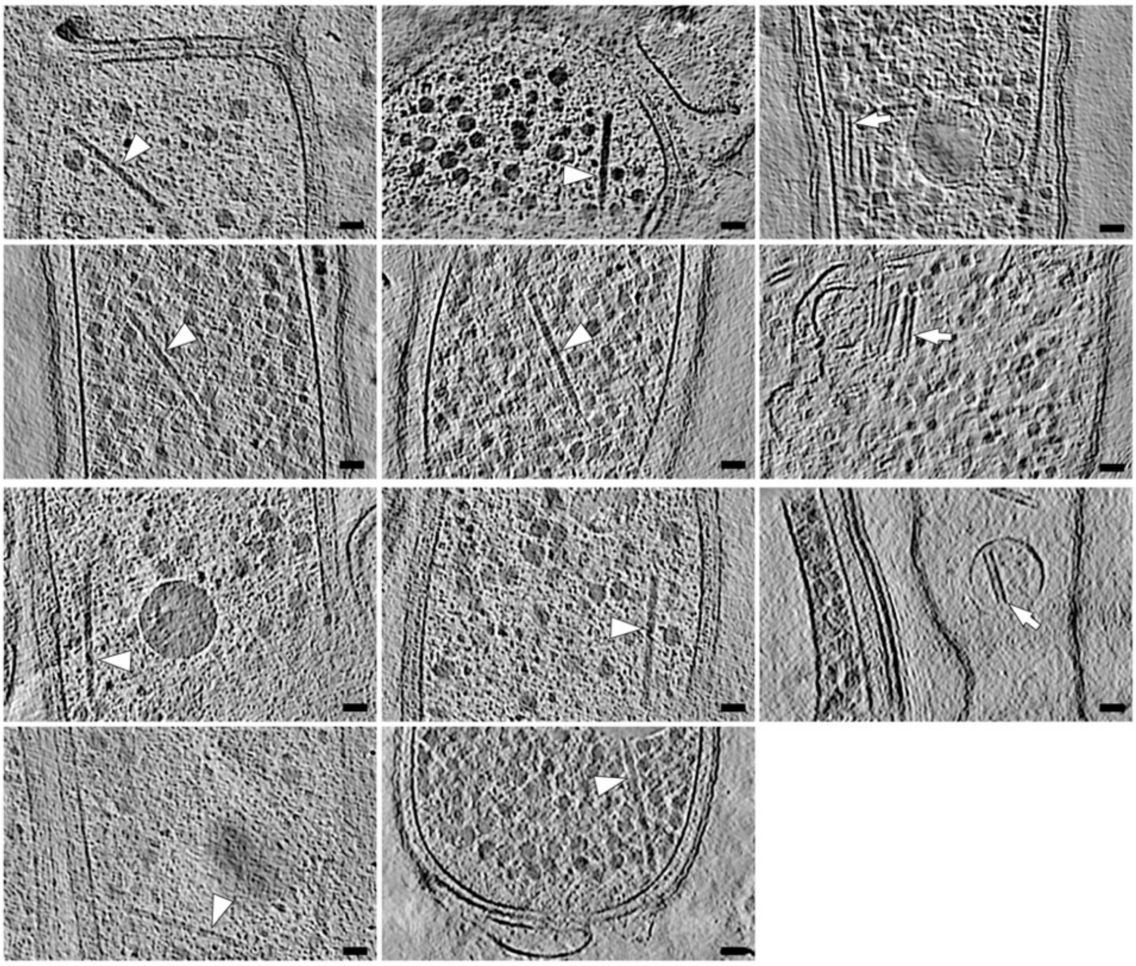

**Figure S5. The CIS-like features are seen in mat *Roseiflexus*-like cells.**
Slices through cryo-tomograms show examples of CIS-like features observed in environmental *Roseiflexus*-like cells from the mat. Arrowheads indicate extended CIS and arrows indicate contracted CIS. Slice thickness: 13.5 and 18 nm. Scale bars: 50 nm.

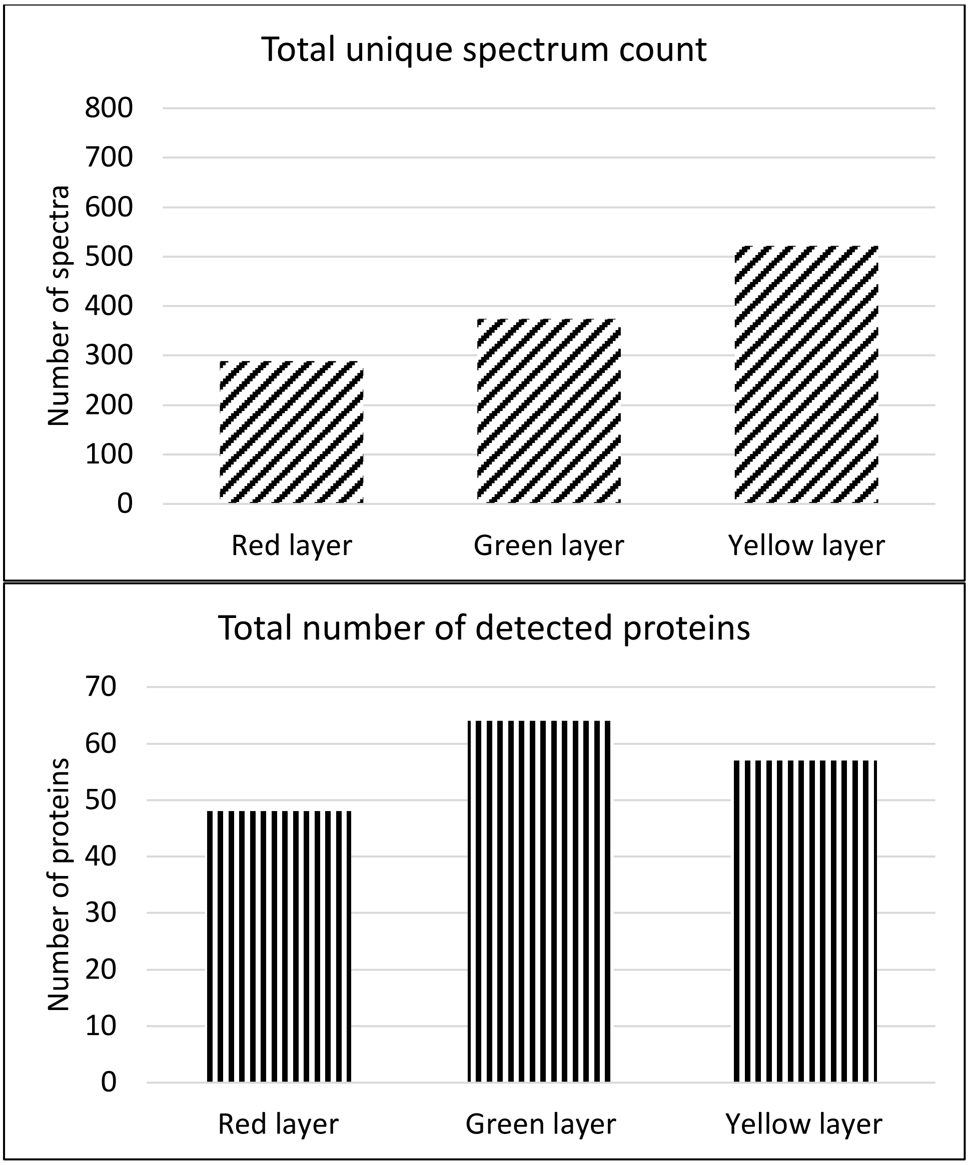


**Figure S6. Mass spectrometry with a search against the whole metagenomes (all contigs longer 1 kb from all layers) detected proteins in the CIS purifications of the different mat layers.** The top bar graph shows result of the total unique spectrum count, bottom bar graph shows total number of deteced proteins.

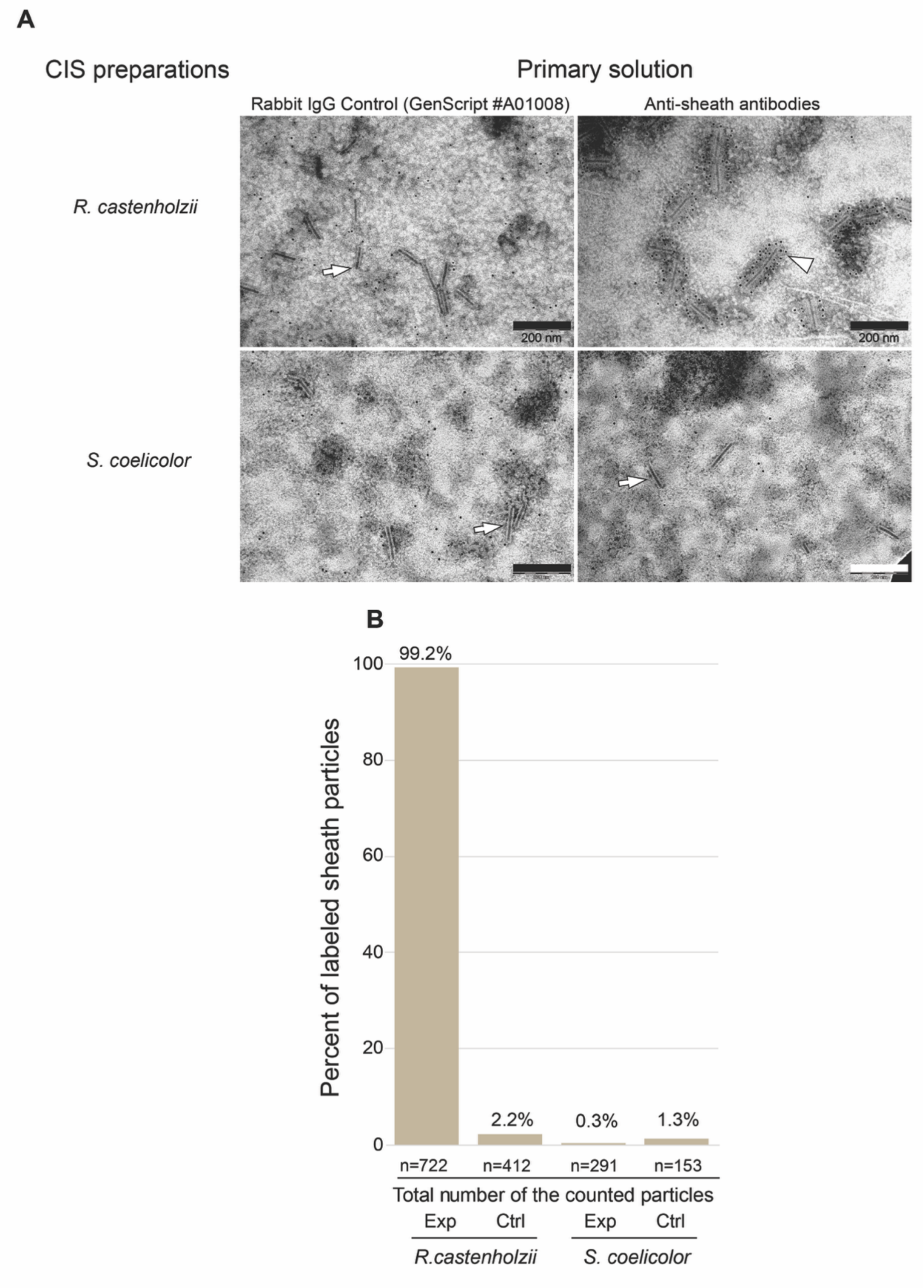


**Figure S7. Application of the rabbit anti-sheath antibodies against *Roseiflexus* sheath protein resulted in specific labelling the CIS particles from *Roseiflexus* cells.** (**A**) Examples of the original TEM microphotographs show labelled (arrowhead) and unlabelled (arrow) CIS particles from *R. castenholzii* and *S. coelicolor* axenic cultures. Scale bars: 200 nm. (**B**) Bar plot showing number of the labelled CIS particles in the CIS preparations from *R. castenholzii* and *S. coelicolor* axenic cultures. Primary anti-sheath rabbit antibodies were applied for labelling the sheath protein (Exp). Rabbit IgG control was applied as control to check the unspecific binding of anti-rabbit gold-conjugated secondary antibodies (Ctrl).

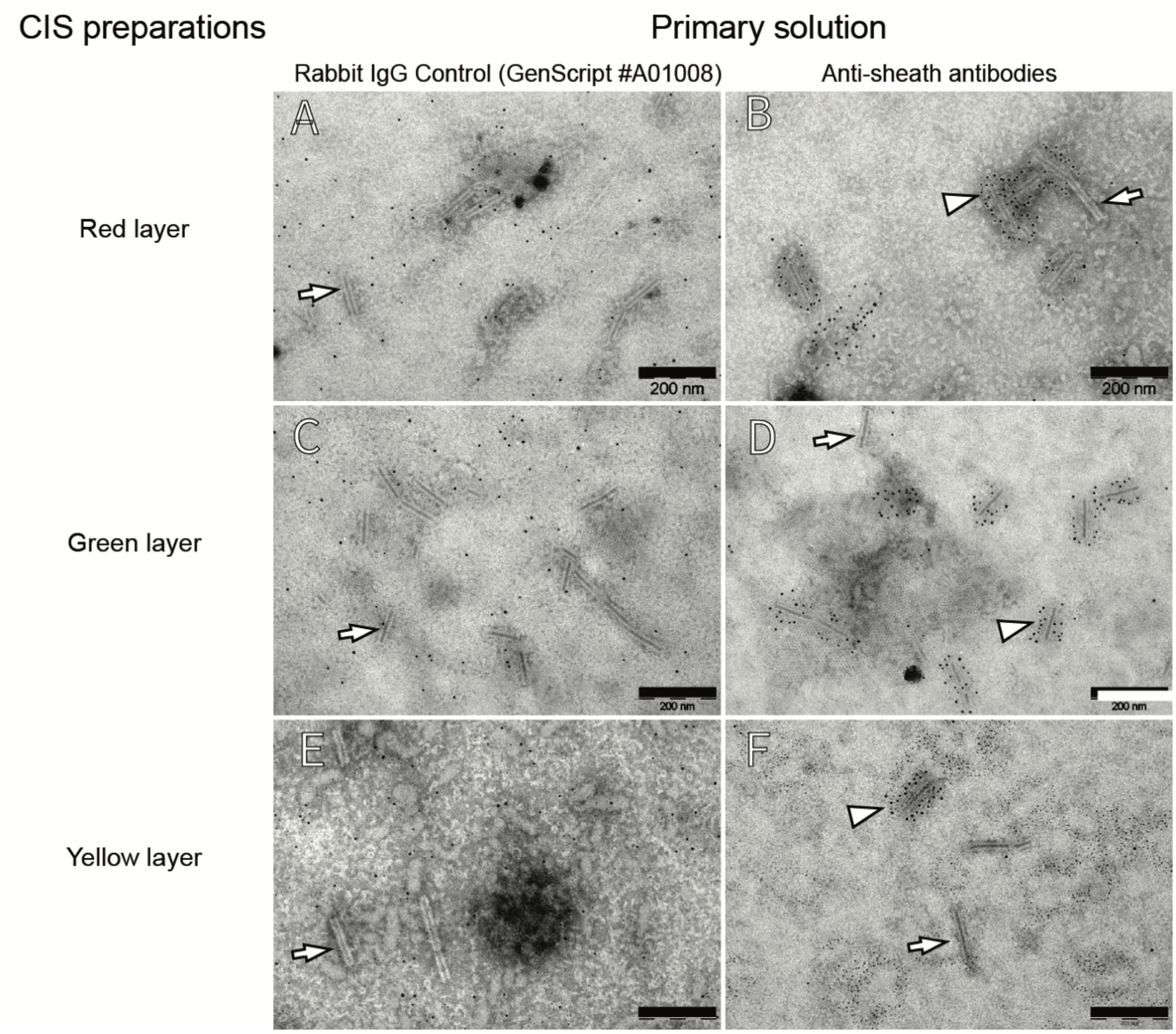


**Figure S8. Application of the rabbit anti-sheath antibodies against *Roseiflexus* sheath protein resulted in specific labelling the CIS particles from the mat.** Examples of the original TEM microphotographs show labelled (arrowhead) and unlabelled (arrow) CIS particles from the mat layers (Red layer – A and B; Green layer – C and D; Yellow layer – E and F). Scale bars: 200 nm.

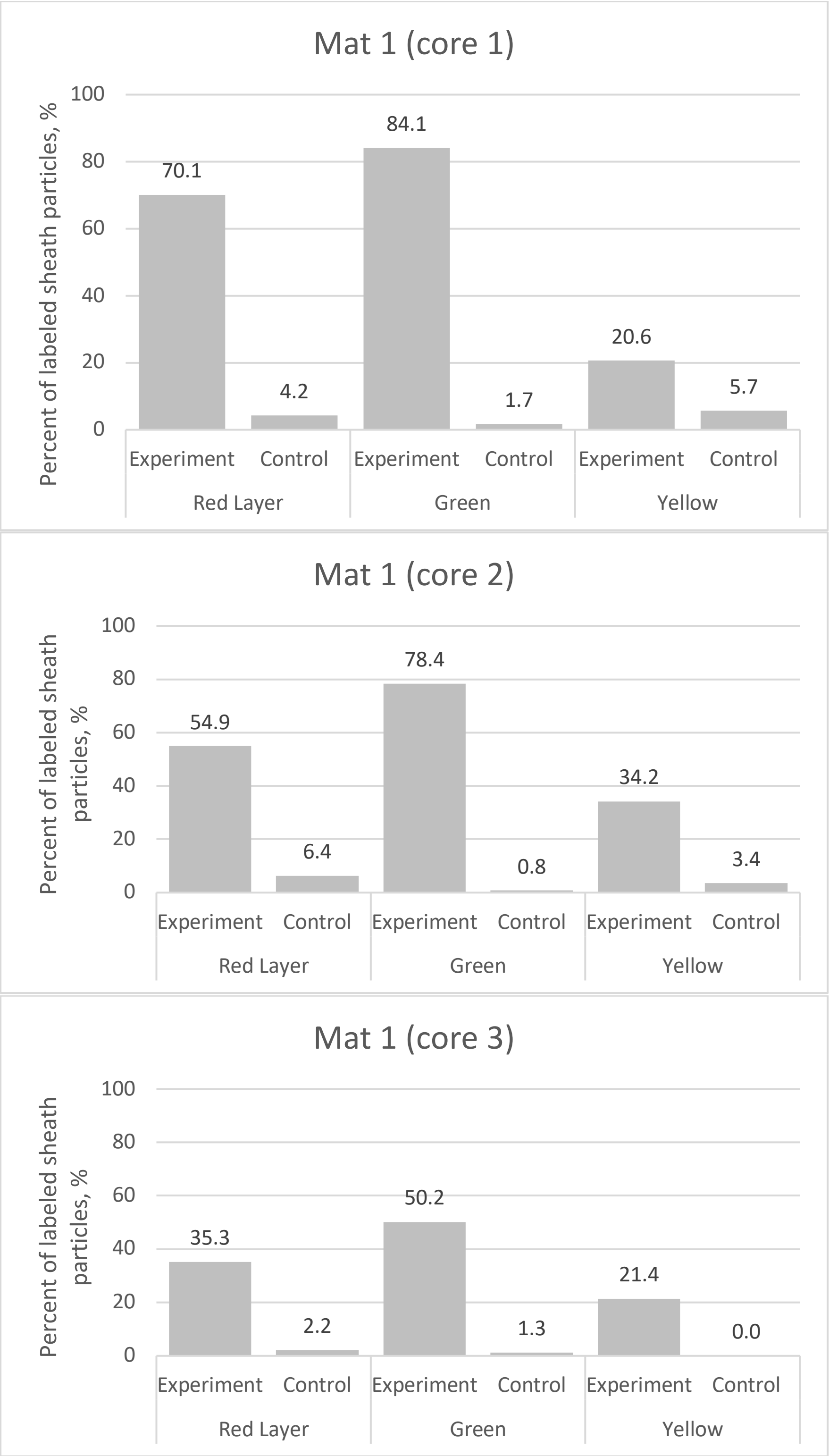


**Figure S9. *Roseiflexus* CIS particles represent a major fraction of the CIS particle pool in preparations from the green layer of the mat according to results of the immunogold labelling.** Bar plots show the percentage of *Roseiflexus* CIS particles in deferent layers of the three mat cores collected from the mat. *Roseiflexus* CIS particles were counted as immunogold-labelled particles in TEM images (see examples of the images in Figure S8).

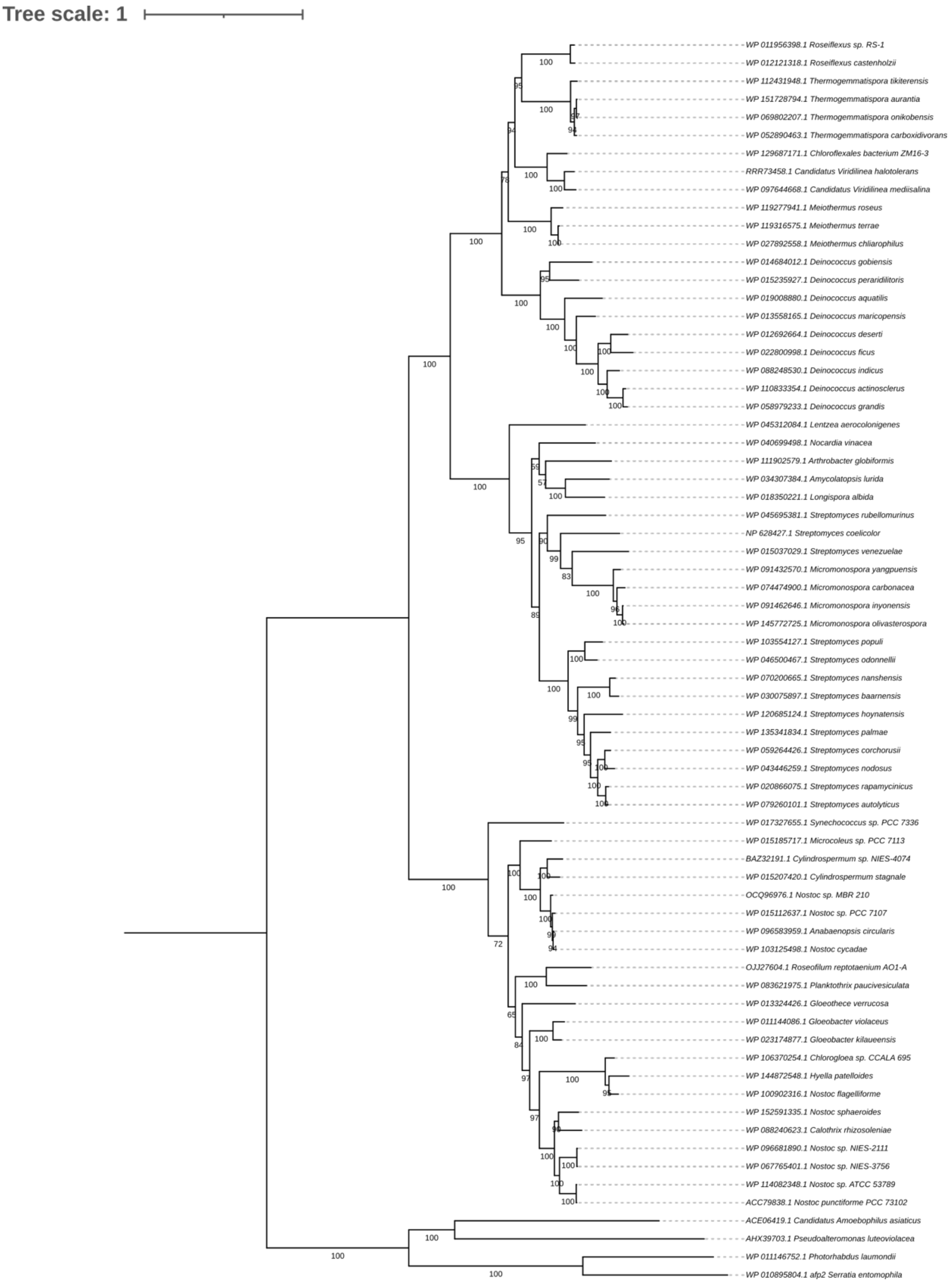


**Figure S10. A maximum likelihood phylogenetic tree shows that a clade of the *Roseiflexus*-related CISs clusters with CISs from *Streptomyces* strains within lineage subtype IId. The tree was reconstructed based on analysis of concatenated sheath and baseplate protein sequences using IQ-TREE 1.6.11 with the LG+F+I+G4 model.**
